# Comparative genomic analysis between *Leonurus japonicus* and *Leonurus sibiricus*

**DOI:** 10.1101/2022.11.27.518111

**Authors:** Dan-Jie Yang, Meng-Xiao Yan, Peng Li, Pan Liu, Yun Gao, Yan Jiang, Ze-Kun He, Yu Kong, Xin Zhong, Sheng Wu, Jun Yang, Hong-Xia Wang, Yan-Bo Huang, Le Wang, Xiao-Ya Chen, Yong-Hong Hu, Qing Zhao, Ping Xu

## Abstract

*Leonurus japonicus* Houtt. is an important medicinal plant in East Asia and is now widely recognized for its role in treating cerebral apoplexy and lowering blood lipids. Here, we report two sets of chromosome-level genome sequences for leonurine-producing *Leonurus japonicus* and for its closely related species leonurine-free *Leonurus sibiricus*, where 99.78% of 518.19 Mb of *L. japonicus* was assembled into ten pseudochromosomes with a contig N50 of 17.62 Mb and 99.33% of 472.29 Mb of *L. sibiricus* was assembled into nine pseudochromosomes with a contig N50 of 13.29 MB. The reference genomes of *Leonurus* will accelerate the decoding of novel bioactive molecules in medicinal plants, especially in the Lamiaceae family.

## Introduction

*Leonurus japonicus* Houtt., or Chinese motherwort, is a member of the family Lamiaceae. It is widely used as traditional medicine for the treatment of gynecological diseases in China, Korea, India, and other Asian countries (Shang et al., 2014). *L. japonicus* is called “Yi Mu Cao” (beneficial herb for women) in China and the name clearly suggests its medicinal properties. The authoritative book on Traditional Chinese Medicine, *Ben Cao Gang Mu* described it as a “panacea in treating diseases about blood” (Miao et al., 2019). Modern pharmacological studies have shown that *L. japonicus* has effects on the uterus as well as cardioprotective (Chen et al., 2017), anti-inflammatory (Li et al., 2020a; Yang et al., 2020; Zhang et al., 2020), hepatoprotective (Tian et al., 2021), antioxidative (Liao et al., 2021), antitumour (Park et al., 2022; Yue et al., 2020), angiogenic (He et al., 2018; Zhou et al., 2020), neuroprotective (Li et al., 2020b) and antibacterial activities (Xiong et al., 2013).

Two alkaloids, leonurine and stachydrine, are the major bioactive compounds that accumulate in *L. japonicus* and mainly account for the pharmaceutical activities of the medicinal plant (Zhang et al., 2018). According to the Pharmacopoeia of China, leonurine and stachydrine are now used as the official indicators to monitor the quality of the herb and of the preparations with *L. japonicus* (Committee for the pharmacopoeia of P.R. China, 2020). Leonurine, also named 4-guanidino-n-butyl syringate, has significant efficacy in the treatment of stroke and lowering blood lipids. This molecule is currently under phase II clinical trials. With the increase in the incidence of stroke worldwide, leonurine has captured the attention of many researchers in medical science. Despite the importance and growing demand for *L. japonicus* and leonurine, the genome of *L. japonicus* has not yet been reported, which limits genetic improvement of the medical plant as well as elucidation of the biosynthetic pathways of its bioactive compounds.

The genomics of medicinal plants have developed rapidly during recent years, which provides useful information to researchers to elucidate the biosynthetic pathways of important natural products, such as *Camptotheca acuminata* and camptothecin (Kang et al., 2021), *Scutellaria baicalensis* and wogonin (Zhao et al., 2019), *Coptis chinensis* and protoberberine-type alkaloids (Liu et al., 2021), *Salvia miltiorrhiza* and tanshinones (Li et al., 2022; Ma et al., 2021), *Tripterygium wilfordii* and triptolide (Tu et al., 2020).

*Leonurus sibiricus* is another species that is closely related to *L. japonicus*. Both are common in the wild in China with similar appearances, but the metabolic patterns are extremely different (Pitschmann et al., 2017; Zhang et al., 2018) because the whole plant of *L. sibiricus* is almost leonurine-free. According to the huge difference in leonurine, *L. japonicus* and *L. sibiricus* are perfect research materials to study the biosynthesis pathway of leonurine. The *de novo* genome sequencing of the two species would offer valuable reference genomic information for *Leonurus* research.

Here, we report chromosome-level genome sequences of *L. japonicus* and *L. sibiricus*, which will greatly accelerate the resolution of the synthesis pathway of active compounds in *Leonurus*, especially leonurine. Understanding the genes responsible for leonurine biosynthesis in plants will lay a foundation for biosynthesis and molecular breeding for improved productivity.

## Results

### Comparative analyses of metabolites between *L. japonicus* and *L. sibiricus*

*L. japonicus* and *L. sibiricus* are two common species found in China and other Asian countries with similar appearances. The karyotype analysis showed that they possessed different chromosome numbers, with *L. japonicus* 2n = 20 and *L. sibiricus* 2n = 18 (Fig. 1A). However, *L. japonicus*, not *L. sibiricus*, has been used as an important medicinal plant by Chinese people for the treatment of gynecological problems for thousands of years, suggesting that there are differences in bioactive ingredients between the two species.

**Fig. 1.**
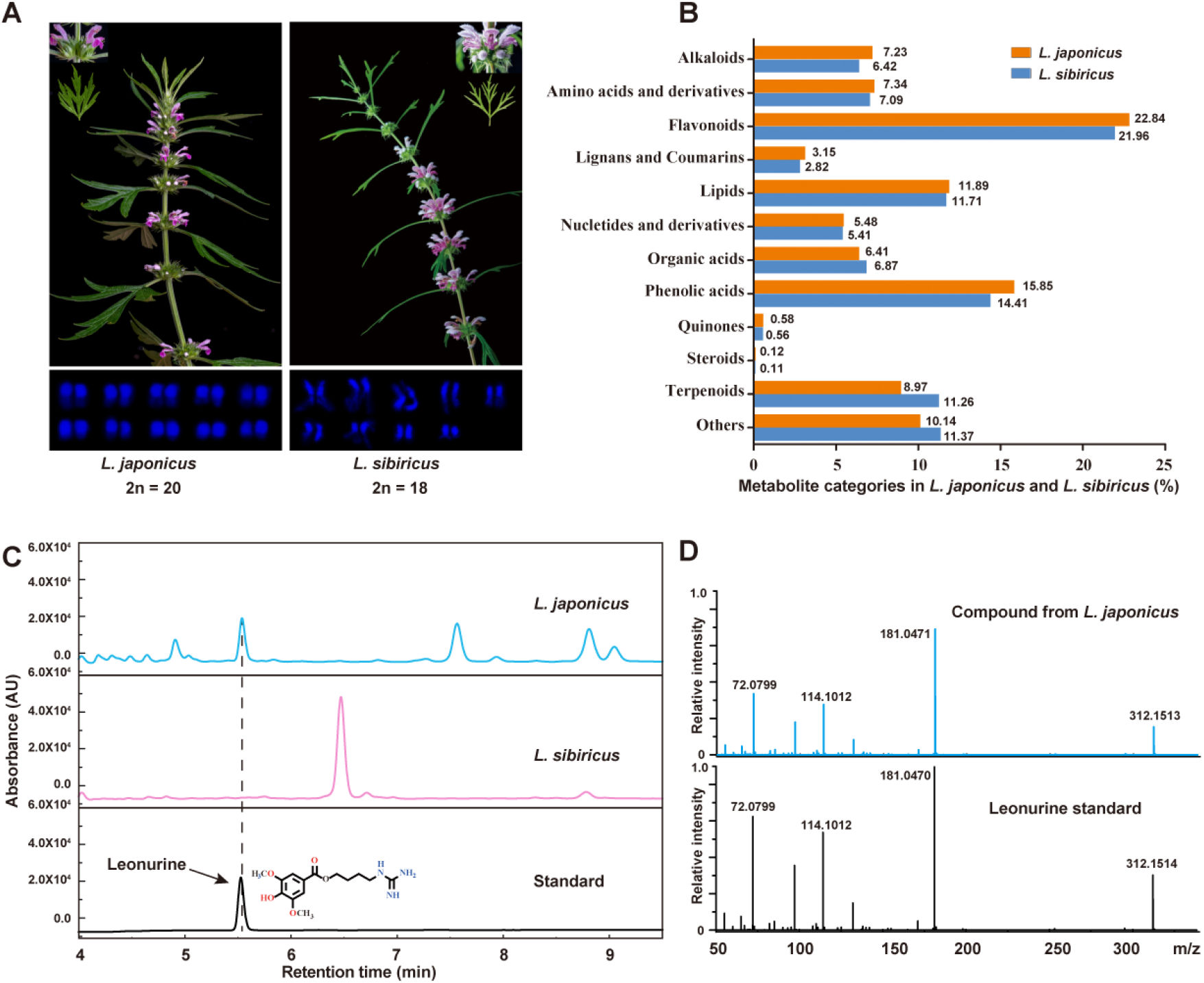
Metabolome analysis of *L. japonicus* and *L. sibiricus*. **A** Morphological differences and karyotype analysis between *L. japonicus* and *L. sibiricus*. **B** Metabolite composition and proportion in *L. japonicus* and *L. sibiricus*. **C** LC-MS analysis of leonurine contents in *L. japonicus* and *L. sibiricus*. **D** MS profiles of the compound from *L. japonicus*, which were identical to the leonurine standard.

Metabolic analysis of the two *Leonurus* species was carried out, and the results showed that *L. japonicus* accumulates leonurine in large amounts in its aerial parts and tiny amounts in roots, but the compound was not detectable in any tissue of *L. sibiricus* (Fig. 1C-D and Fig. S1). A broad-target metabolome analysis returned a total of 858 substances in *L. japonicus* and 888 substances in *L. sibiricus* (Fig. S2). The results showed that the categories of alkaloids, amino acid derivatives and flavonoids in *L. japonicus* were slightly more abundant than those in *L. sibiricus*, while terpenoids were more diversified in *L. sibiricus* (Fig. 1B).

Together, these results reveal the major differences in metabolites and active ingredients between the two species and confirm that leonurine is only synthesized in *L. japonicus*, which prompted us to further explore comparative genomics between the two species.

### High-quality sequencing and assembly of two *Leonurus* genomes

We sequenced and assembled the two *Leonurus* genomes using Oxford Nanopore Technologies (ONT) and improved the accuracy of the assembly using sequencing data obtained from the Illumina platform. An overview of the sequencing data is shown in Table 1. Based on the *k*-mer distribution analysis, we estimated a genome size of 553.75 Mb for the *L. japonicus* genome with a heterozygosity of 0.73% and a genome size of 465.78 Mb for the *L. sibiricus* genome with a heterozygosity of 1.91% (Fig. S3, Table S1 and S2). In total, we generated 55.75 Gb and 34.14 Gb of data for *L. japonicus* and *L. sibiricus*, representing approximately 100× and 73× coverage of the predicted genomes, respectively.

**Table 1.** The statistics of the assemblies and annotations of two *Leonurus* genomes.

|  | *L. japonicus* | *L. sibiricus* |
| --- | --- | --- |
| **Assembly** |  |  |
| Total number of contigs | 73 | 74 |
| Assembly size (MB) | 518.19 | 472.29 |
| Contig N50 (MB) | 17.62 | 13.29 |
| Contig N90 (MB) | 5.74 | 3.33 |
| Largest contig (MB) | 50.97 | 37.86 |
| Total number of scaffolds | 15 | 15 |
| Assembly size (MB) | 518.2 | 472.3 |
| Scaffold N50 (MB) | 53.88 | 55.54 |
| Scaffold N90 (MB) | 43.56 | 40.86 |
| Largest scaffold (MB) | 61.92 | 71.62 |
| anchored rate | 99.78% | 99.33% |
| DNA mapping rate (Illumina reads) | 99.94% | 99.55% |
| BUSCO | 97.90% | 96.41% |
| Total GC content | 35.29% | 35.62 |
| **Annotation** |  |  |
| Protein-coding genes | 28664 | 25748 |
| Repetitive sequences | 66.51% | 65.39% |
| Number of ncRNA | 4576 | 3955 |
| BUSCO | 98.40% | 96.90% |

The total length of the *L. japonicus* genome assembly was 518.19 Mb with a contig N50 value of 17.62 Mb and 472.29 Mb, with a contig N50 of 13.29 Mb for *L. sibiricus*. Benchmarking Universal Single Copy Orthologues (BUSCO) analysis indicated that genome assembly completeness was 97.9% for *L. japonicus* and 96.41% for *L. sibiricus*. The *L. japonicus* genome had a 99.94% DNA mapping rate, and for *L. sibiricus*, the DNA mapping rate was 99.55%, suggesting that the two genome assemblies are high-quality. We further connected the contigs into pseudochromosomes using a Hi-C matrix. In total, 99.78% and 99.33% of the assembled *L. japonicus* and *L. sibiricus* contigs were correctly anchored into ten and nine pseudochromosomes, respectively, with N50 values of 53.88 Mb and 55.54 Mb, respectively. The strong signal along the diagonal of interactions between proximal regions suggested the high quality of the Hi-C assemblies for the *L. japonicus* and *L.sibiricus* genomes (Fig. S4). Hence, two sets of high-quality assemblies and chromosome-level genome sequences are provided here, providing a valuable source for the study of *Leonurus* and other Lamiaceae species.

### Repeat and gene annotations

Using the combination of *ab initio* and homology-based analysis, together with RNA-Seq read-assisted annotation, we annotated 28,664 and 25,748 protein-coding genes in the genomes of *L. japonicus* and *L. sibiricus*, respectively; 93.69% of the *L. japonicus* and 94.76% of the *L. sibiricus* genes had functional annotations. The average transcript sizes were 4,432.94 bp in *L. japonicus* and 4,429.48 bp in *L. sibiricus*. In addition, 344.64 Mb and 308.85 Mb of repetitive elements were identified by the combination of ab initio and homology-based approaches, occupying approximately 66.51% and 65.39% of the *L. japonicus* and *L. sibiricus* genomes, respectively. For the two major members of long terminal repeats (LTRs), LTR/Copia and LTR/Gypsy accounted for 41.1% and 40.67% of the genomes of *L. japonicus* and *L. sibiricus*, respectively (Table S3 and S4). A total of 349,492 and 303,570 SSRs were annotated in *L. japonicus* and *L. sibiricus*, respectively, which will provide useful molecular markers for breeding and genetic diversity studies. The *L. japonicus* genome contains 4576 noncoding RNAs (ncRNAs), and for *L. sibiricus*, 3955 ncRNAs were annotated (Table S5 and S6). Consistent with the genome assembly quality assessment, the BUSCO indexes were 98.4% in *L. japonicus* and 96.9% in *L. sibiricus*, suggesting the high completeness of the genome annotations.

### Comparative genomic analysis between *L. japonicus* and *L. sibiricus*

Comparative genomic analyses were carried out between the two *Leonurus* species based on their high-quality genomes. *L. japonicus* has one more pair of chromosomes than *L. sibiricus*, and the overall comparison of the genomes is shown in Fig. 2A-B. The collinearity analysis of the two genomes revealed that chromosome breakage and rearrangement resulted in one more pair of chromosomes in *L. japonicus*. Chromosome 2 of *L. japonicus* was derived from the breakage and fusion of chromosomes 2, 3 and 4 of *L. sibiricus*, chromosome 3 was derived from chromosomes 3 and 4 of *L. sibiricus*, and chromosomes 1 and 3 of *L. sibiricus* together formed chromosome 5 of *L. japonicus* (Table S7). Despite the structural variation, there is a large degree of collinearity between the genomes of the two *Leonurus* species, and the vast majority of genes are colinear between the two species.

**Fig. 2.**
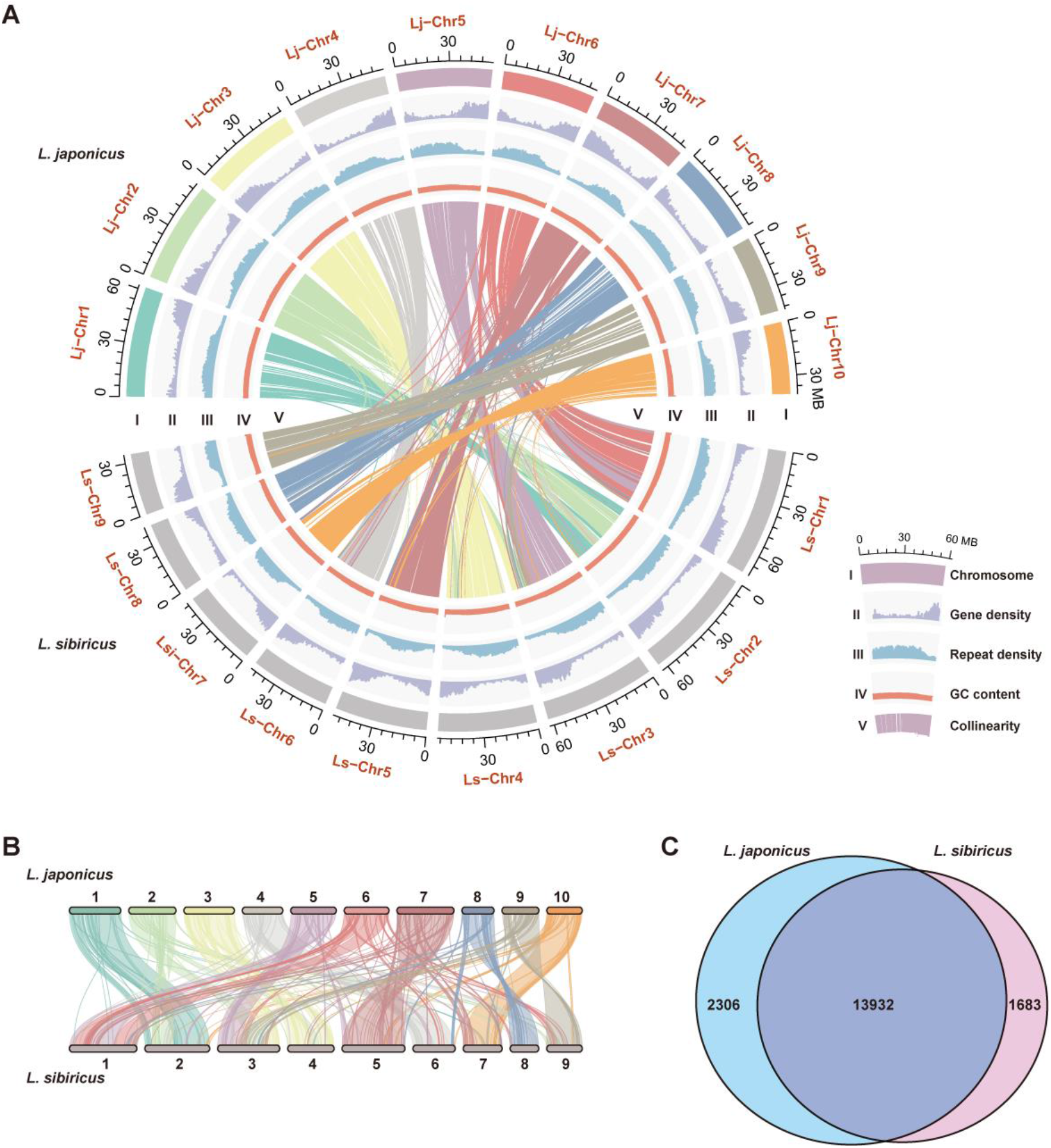
Genome analysis of *L. japonicus* and *L. sibiricus*. **A** An overview of genome assemblies and annotations of *L. japonicus* and *L. sibiricus*. (I) Circular representation of the pseudomolecule. (II–IV) Gene density (50 kb window), percentage of repeats (50 kb window), and GC content (50 kb window). (V) Each linking line in the center of the circle connects a pair of homologous genes. **B** Collinear relationship between *L. japonicus* and *L. sibiricus* chromosomes. **C** Shared and unique gene families in the *L. japonicus* and *L. sibiricus* genomes.

Analysis of gene families also revealed a large degree of overlap between the two gene families, with approximately 86% of the gene families being common to both species (Fig. 2C).

### Phylogenetic and whole-genome duplication analyses

To investigate the evolutionary history of two *Leonurus* species, we performed phylogenetic analysis with 12 other sequenced genomes (*Oryza sativa, Arabidopsis thaliana, Scutellaria baicalensis, Salvia miltiorrhiza, Sesamum indicum, Catharanthus roseus, Solanum lycopersicum, Amborella trichopoda, Nicotiana attenuata, Camptotheca acuminata, Abrus precatorius and Papaver somniferum*). The five Lamiales species formed a monophyletic clade, including four Lamiaceae species (*L. japonicus, L. sibiricus, Scu. baicalensis* and *Sal. miltiorrhiza*) and a Pedaliaceae species (*Ses. indicum*), with a divergence time of approximately 42.4 MYA, which was consistent with previous reports (Xu et al., 2020). *L. japonicus* and *L. sibiricus* formed a small monophyletic clade, as the recent diverged clade in the Lamiales clade, with an estimated divergence time of approximately 3.8 MYA (Fig. 3A), following the divergence of their common ancestor from *Scu. baicalensis* at approximately 26 MYA. The collinearity analysis of two *Leonurus* species with grape indicated that they each performed a WGD event in addition to whole-genome triplication (γ event). To further determine whether a WGD occurred in Lamiales species, we performed a curve-fitting analysis of the synonymous substitution rate (Ks) distribution of homologous genes (Fig. 3B). The results showed that there were two sharp duplication peaks with close positions, which was consistent with the collinearity analysis results, suggesting that the five Lamiales species underwent another WGD event after γ event.

**Fig. 3.**
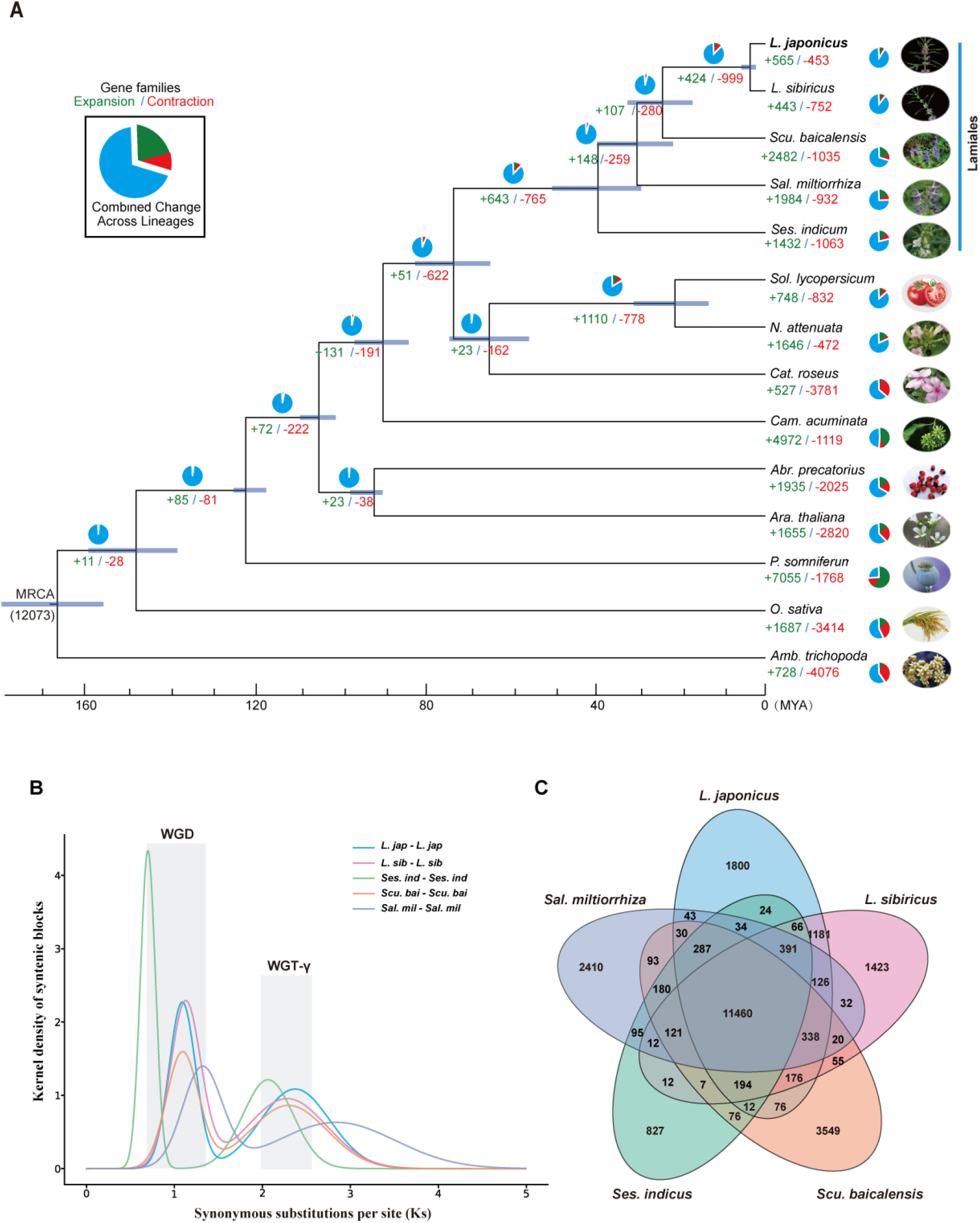
Phylogenetic and whole-genome duplication analyses. **A** Phylogenetic tree showing the topology and divergence times for 14 plant species. The blue bars show the 95% confidence interval of divergence time, and numbers in green and red at branches indicate the expansion and contraction of gene families, respectively. The pie chart colors represent the changes in the gene family (blue, no significant change; green, expanded; red, contracted). **B** Synonymous substitution rate (Ks) distributions of syntenic blocks for the paralogues and orthologues of *Scu. baicalensis, Sal. miltiorrhiza, Ses. indicum, L. japonicus* and *L. sibiricus*. **C** The Venn diagram shows the sharing and specificity of gene families presented among five Lamiales species.

The gene families of the five Labmiales species are highly conserved, since the majority of gene families (11460) are shared, and only 827-3546 gene families are unique to each species (Fig. 3C). Compared with their most recent common ancestor (MRCA), 565 and 443 gene families were expanded in *L. japonicus* and *L. sibiricus* lineages, respectively, while 453 and 752 gene families were contracted, respectively, which are significantly less than the numbers of expansions and contractions in other species of Labmiales (Fig. 3A). Gene Ontology (GO) enrichment analyses of the expanded gene families in *L. japonicus* and *L. sibiricus* showed that they were significantly enriched in “monooxygenase activity”, “serine-type carboxypeptidase activity”, ‘‘alcohol O-acetyltransferase activity’’, “acyltransferase activity’’ and “sinapine esterase activity”, which are likely involved in the biosynthesis of abundant secondary metabolites in the two *Leonurus* species (Fig. S5A-B and Fig. 3B). Among them, the expansion of “serine-type carboxypeptidase activity” specifically occurred in *L. japonicus* and may be associated with leonurine biosynthesis (Fig. S5A-B).

## Discussion

The Lamiaceae family contains a variety of valuable medicinal plants, such as *Scu. baicalensis* and *Sal. miltiorrhiza*. Lamiaceae plants produce various secondary metabolites, and most of them are important for plants to adapt to the natural environment and provide flavoured foods and natural medicines for human health (Mint Evolutionary Genomics Consortium. Electronic address and Mint Evolutionary Genomics, 2018). Plant secondary metabolites are complex, diverse in structure, and have species specificity, which makes them difficult to study. In recent years, the advancement of sequencing technology and the reduction of sequencing costs, combined with the improvement of metabolomic detection databases, have led to the rapid development of medicinal plant research. The genome of medicinal plants provides an extremely useful tool for evolutionary analysis of the synthetic pathway of bioactive substances. Motherwort is a traditional medicinal plant with a long history of treating gynecological ailments (Zhang et al., 2018). In recent years, studies on motherwort have made much progress in the treatment of neurological diseases and reducing blood lipids, which has been widely recognized (Jin et al., 2019; Law et al., 2017; Meng et al., 2019; Zhang et al., 2018; Zhu et al., 2018). At present, leonurine, derived from motherwort, is being tested in clinical trials as a new class of drug for the treatment of stroke and lowering blood lipids in China.

*Leonurus* is a relatively small but very distinctive genus. There are approximately 24 species worldwide. A large number of studies show that leonurine is the main active ingredient of *Leonurus* and has a variety of beneficial values. We obtained two species of *Leonurus* with a tremendous difference in leonurine content and obtained high-quality genomes at the chromosome level of both species, which provided a basis to analyse the synthetic pathway of leonurine in plants. At the same time, it is also of great significance to analyse the evolution of metabolites in the family Lamiaceae.

## Methods and materials

### Plant Materials, DNA library construction and sequencing

Fresh leaves and roots were collected from *L. japonicus* and *L. sibiricus* grown in Shanghai Chenshan Botanical Garden. High-quality genomic DNA was isolated from the fresh leaves of *L. japonicus* and *L. sibiricus* using the CTAB method, and total mRNA was extracted using the RNA Prep Pure Plant Plus Kit according to the manufacturer’s instructions (Tiangen Biotech, Beijing, China). A strand-switching method and the cDNA-PCR Sequencing Kit (SQK-PCS109) were used to reverse transcribe mRNA into cDNA. The DNA quality and concentration were tested by 0.75% agarose gel electrophoresis, a NanoDrop One spectrophotometer (Thermo Fisher Scientific) and Qubit 3.0 Fluorometer (Life Technologies, Carlsbad, CA, USA).

After DNA samples were tested and passed, they were randomly sheared by a Covaris ultrasonic disruptor. Illumina sequencing pair-end libraries with an insert size of 300 bp were prepared using the Nextera DNA Flex Library Prep Kit (Illumina, San Diego, CA, USA). After the libraries were constructed, Qubit 2.0 was used for initial quantification, and Agilent 2100 was used for quality control of the libraries after the libraries were diluted and the insert sizes were as expected. Sequencing was subsequently performed using the Illumina NovaSeq6000 platform (Illumina, San Diego, CA, USA). The SOAPnuke (v2.1.4) tool (https://github.com/BGI-flexlab/SOAPnuke) was used to filter raw reads and obtain clean reads for genome survey analysis and genome polishing. For long-read DNA sequencing, the libraries were prepared using the SQK-LSK109 ligation kit. The purified library was sequenced using a PromethION sequencer (Oxford Nanopore Technologies, Oxford, UK) at Wuhan Benagen Tech Solutions Company Limited (Wuhan, China).

### Genome assembly and assessment

The GCE software (version 1.0.0) was used to estimate the genome size of *L. japonicus* and *L. sibiricus*. The heterozygosity and repeat rate of the two *Leonurus* genomes were determined using the *k*-mer method with Jellyfish software (*http://www.genome.umd.edu/jellyfish.html*). The preliminary genome assemblies were first performed using nextDenovo software (https://github.com/Nextomics/NextDenovo). Then, two rounds of error correction were performed on the assembly results based on the nanopore sequencing data using Racon (v1.4.11) (https://github.com/isovic/racon) and Pilon (v1.23) was used to further correct twice based on the Illumina Novaseq sequencing data. Finally, the genome was removed from the heterozygous sequences using the Purge_haplotigs pipeline (v.1.0.4) to obtain the final assembly result. The consistency and completeness of the two assembled genomes were evaluated by mapping rate, average depth of the Illumina short reads and BUSCO software (v.4.1.2).

### Hi-C sequencing and data processing

High-quality DNA extracted from young leaves of *L. japonicus* and *L. sibiricus* was used for Hi-C sequencing. Formaldehyde was used for fixing chromatin. In *situ* Hi-C chromosome conformation capture was performed according to the DNase-based protocol described by Ramani. The libraries were sequenced using 150 bp paired-end mode on an Illumina NovaSeq (Illumina, San Diego, CA, USA). For pseudochromosome-level scaffolding, we used the assembly software ALLHIC (v0.9.12) for stitching, and then we imported the final files (.hic and.assembly) generated by the software into Juicebox (v1.11.08) for manual optimization.

### Gene prediction

Evidence from transcript mapping, ab initio gene prediction, and homologous gene alignment was combined to predict protein-coding genes in the *L. japonicus* and *L. sibiricus* genomes. ONT cDNA reads from *L. japonicus* and *L. sibiricus* were aligned against the genome using Minimap2 (v2.17). Transcripts were assembled using stringtie2 (v2.1.5), and all assembled transcript ORFs were predicted by Trans-Decoder (v5.1.0) (https://github.com/TransDecoder/TransDecoder). Augustus (v3.3.2), Genscan (v1.0) (http://bioinf.uni-greifswald.de/webAugustus/predictiontutorial) and GlimmerHMM (v3.0.4) (http://ccb.jhu.edu/software/glimmer/index.shtml) were used for ab initio gene prediction. For homologous gene alignment, the proteins from four related species (*L. japonicus, L. sibiricus, Scu. baicalensis, Sal. miltiorrhiza* and *Ses. indicum*) were aligned to the genome using Exonerate (v2.4.0) (https://github.com/nathanweeks/exonerate). Finally, MAKER (v2.31.10) software (http://yandell.topaz.genetics.utah.edu/cgi-bin/maker_license.cgi) was used to integrate gene sets predicted by the three methods and remove incomplete genes and genes with too short a CDS (CDS length < 150 bp), and a nonredundant and more complete gene set was obtained. BUSCO software (v.5.2.2) was employed (Manni et al. 2021) to evaluate the quality of the prediction.

### Gene function annotation

Functional annotation of the predicted protein-coding genes was carried out by performing Blastp searches with cut-off e-values of 1e-5 against entries in both the NCBI nr and UniProt databases (http://www.UniProt.org/). Searches for gene motifs and domains were performed using InterProScan (v5.33) and HMMER (v3.1). The GO terms (http://geneontology.org/) for genes were obtained from the corresponding InterPro (https://github.com/ebi-pf-team/interproscan) or UniProt entry (https://www.uniprot.org/). Pathway annotation was performed using KOBAS (v3.0) (https://github.com/xmao/kobas) against the KEGG database.

### Comparative genomics for phylogenomic analysis and gene family

To investigate the evolutionary history of the two *Leonurus* genomes, the relationships of these protein-coding genes with those of 12 other angiosperms (*O. sativa, Ara. thaliana, Scu. baicalensis, Sal. miltiorrhiza, Ses. indicum, Cat. roseus, Sol lycopersicum, Amb. trichopoda, N. attenuata, Cam. acuminata, Abr. precatorius* and *P. somniferum*) were aligned using Blastp (v.2.6.0) with cut-off e-values of 1e-5, and gene family clustering was performed using OrthoMCL software (v.2.0.9) (percent Match Cut-off = 30, e-value Exponent cut-off = 1e-5, expansion coefficient =1.5). A total of 246 single-copy gene families shared by 14 selected species were screened to construct phylogenetic trees. First, MUSCLE (v.3.8.31) software was used to align the predicted proteins of single-copy genes, and the TRIMAL (v.1.4.rev22) program (http://trimal.cgenomics.org/introduction) was used to filter the comparison results (-gt0.2). Gene trees were then reconstructed for each gene family using RAxML (v.8.2.10) software with 100 replicates of bootstrapping. For each gene family, we estimated four different gene trees based on amino acid alignments, DNA alignment, codon alignment (nucleotides forced to the amino acid alignment), and codon 1 and 2 alignment (codon alignments where the third codon position was removed). Nucleotide-based analyses were conducted using the GTR + GAMMA model; for amino acid analyses, the WAG model was used. For four different datasets, four maximum likelihood trees of species were inferred from gene trees using RAxML (v.8.2.10) software. A phylogenetic tree of each single-copy gene was further constructed to infer a consensus species tree using ASTRAL (v5.7.1).

Based on the phylogenetic tree results and published divergence times on TIMETREE (http://www.timetree.org/), *Amb. trichopoda* and *O. sativa*: 168-194 MYA; *O. sativa* and *P. somniferum*: 148-173 MYA; *P. somniferum* and *Ara. thaliana* 122-134 MYA; *Ara. thaliana* and *Abr. precatorius*: 97-109 MYA, the MCMCTree of PAML (v.4.9) (n sample = 1, 000, 000; burnin = 200, 000; seq type = 0; clock = 3; model = 4) was used to estimate the divergence time of the different species.

Gene family contraction and expansion analysis in the 14 species was performed using CAFE (v.2.1) software based on gene family clustering results from OrthoMCL analysis; this software employed a random birth and death model to estimate the size of each gene family.

### Whole-gene duplication analysis

KS-based age distributions for all the paralogues of *L. japonicus* or *L. sibiricus* were constructed using WGDI (Sun et al., 2022) software. All potential paralogues were detected with all-vs-all protein sequence blast using BLASTP with an E-value threshold of e^−5^. First, five related species (*L. japonicus, L. sibiricus, Scu. baicalensis, Sal. miltiorrhiza* and *Ses. indicum*) amino acid and CDSs were self-aligned using Blast with an E-value threshold of e^−5^. Second, all syntenic blocks were identified using the improved collinearity pipeline in WGDI with “*p* value = 0.2”. Then, the Ks value for each anchor gene pair located in syntenic blocks was calculated using the Ks pipeline in WGDI. Next, a Ks dotplot of all anchor pairs was obtained using the block pipeline in WGDI, and the Ks peak pipeline in WGDI was used for distribution analysis of the Ks median value for each syntenic block. Finally, the Ks distribution curve was obtained by Gaussian fitting.

### Extraction of leonurine and other metabolites

Plant materials were collected and frozen in liquid nitrogen immediately, then finely ground and stored at -80°C. The samples were freeze-dried in a freeze dryer (Thermo Scientific) at -50℃ for 48 h before extraction. An amount of 10 mg dry sample was weighed and extracted with 2 mL 70% methanol (v/v) in an ultrasonic bath of ice-water mixture (PL-S40T, Kangshijie, China) for 2 h. The samples were then placed at 4°C overnight for extraction. The extracts were centrifuged at 12,000 × g for 10 min at 4°C (Eppendorf, Germany), and the supernatants were collected. The supernatant was filtered through a 0.22 μm PES syringe filter (Anpel, Shanghai) before detection.

### Qualitative metabolite detection

Leonurine was determined using a coupled Dionex UltiMate 3000 HPLC system, and a Q Exactive Plus Mass Spectrometer (Thermo Scientific) collected the MS (mass spectrum) data in positive-ion mode with a spray voltage of 3.5 kV and a capillary temperature of 320°C. The stepped normalized collision energy (NCE) was set at 30, 50 and 80. Separation was carried out on a 100 × 2 mm 3 μm Luna C18 (2) column. Mobile phases with H_2_O consisting of 0.1% formic acid (A) versus acetonitrile containing 0.1% formic acid (B) were used. The gradient profile was performed as follows: 0 -10 min: 5% - 40% B; 10 -15 min: 40% - 90% B; 15 - 20 min: 90%-98% B; 20 - 21 min: 98% - 5% B. The temperature of the column was maintained at 40 ℃. The flow rate of the mobile phase was 0.4 mL/min with a 280 nm detection wavelength.

### Standard compounds

All chemicals were analytical grade. Leonurine was purchased from Yuanye-Biotech (Shanghai, China) and was dissolved in dimethyl sulfoxide (DMSO).

## Acknowledgements

This work was supported by the National Natural Science Foundation of China (32170349) and the Chenshan Special Fund for Shanghai Landscaping Administration Bureau Program (G232402) to PX.

## Author Contributions

PX initiated the program and coordinated the project. PX, QZ, D-J Y, PL and M-X Y wrote the manuscript. YJ, PL, PL, D-J Y, YG, Y-B H and Z-K H maintained and prepared the samples. PX, M-X Y, Y-H H, and X-Y C designed the sequencing strategy and performed sequencing. YJ, PL, YK and YG performed metabolite measurements. XZ took the photos of plants. D-J Y, M-X Y, YG, LW, H-X W and JY performed the data analysis.

## Competing financial interests

The authors declare no competing financial interests.

## Notes

### Competing Interest Statement

The authors have declared no competing interest.

## References

Chen, H.H., Zhao, P., Zhao, W.X., Tian, J., Guo, W., Xu, M., Zhang, C., and Lu, R. (2017). Stachydrine ameliorates pressure overload-induced diastolic heart failure by suppressing myocardial fibrosis. American journal of translational research 9:4250–4260.

Fu, R., Zhang, P., Jin, G., Wang, L., Qi, S., Cao, Y., Martin, C., and Zhang, Y. (2021). Versatility in acyltransferase activity completes chicoric acid biosynthesis in purple coneflower. Nat Commun 12:1563.

He, Y.L., Shi, J.Y., Peng, C., Hu, L.J., Liu, J., Zhou, Q.M., Guo, L., and Xiong, L. (2018). Angiogenic effect of motherwort (Leonurus japonicus) alkaloids and toxicity of motherwort essential oil on zebrafish embryos. Fitoterapia 128:36–42.

Hong, Z.Y., Yu, S.S., Wang, Z.J., and Zhu, Y.Z. (2015). SCM-198 Ameliorates Cognitive Deficits, Promotes Neuronal Survival and Enhances CREB/BDNF/TrkB Signaling without Affecting Aβ Burden in AβPP/PS1 Mice. International journal of molecular sciences 16:18544–18563.

Jiang, T., Ren, K., Chen, Q., Li, H., Yao, R., Hu, H., Lv, Y.C., and Zhao, G.J. (2017). Leonurine Prevents Atherosclerosis Via Promoting the Expression of ABCA1 and ABCG1 in a Pparγ/Lxrα Signaling Pathway-Dependent Manner. Cellular physiology and biochemistry : international journal of experimental cellular physiology, biochemistry, and pharmacology 43:1703–1717.

Jin, M., Li, Q., Gu, Y., Wan, B., Huang, J., Xu, X., Huang, R., and Zhang, Y. (2019). Leonurine suppresses neuroinflammation through promoting oligodendrocyte maturation. J Cell Mol Med 23:1470–1485.

Kang, M., Fu, R., Zhang, P., Lou, S., Yang, X., Chen, Y., Ma, T., Zhang, Y., Xi, Z., and Liu, J. (2021). A chromosome-level Camptotheca acuminata genome assembly provides insights into the evolutionary origin of camptothecin biosynthesis. Nat Commun 12:3531.

Law, B.Y.K., Wu, A.G., Wang, M.J., and Zhu, Y.Z. (2017). Chinese Medicine: A Hope for Neurodegenerative Diseases? J Alzheimers Dis 60:S151–S160.

Li, C.Y., Yang, L., Liu, Y., Xu, Z.G., Gao, J., Huang, Y.B., Xu, J.J., Fan, H., Kong, Y., Wei, Y.K., et al. (2022). The sage genome provides insight into the evolutionary dynamics of diterpene biosynthesis gene cluster in plants. Cell Rep 40:111236.

Li, H.Y., Li, Y., Wei, W.J., Ma, K.L., Chen, J.J., and Gao, K. (2020a). Halimane and labdane diterpenoids from Leonurus japonicus and their anti-inflammatory activity. Phytochemistry 172:112280.

Li, L., Sun, L., Qiu, Y., Zhu, W., Hu, K., and Mao, J. (2020b). Protective Effect of Stachydrine Against Cerebral Ischemia-Reperfusion Injury by Reducing Inflammation and Apoptosis Through P65 and JAK2/STAT3 Signaling Pathway. Frontiers in pharmacology 11:64.

Liao, L., Gong, L., Zhou, M., Xue, X., Li, Y., and Peng, C. (2021). Leonurine Ameliorates Oxidative Stress and Insufficient Angiogenesis by Regulating the PI3K/Akt-eNOS Signaling Pathway in H(2)O(2)-Induced HUVECs. Oxidative medicine and cellular longevity 2021:9919466.

Lin, M., Pan, C., Xu, W., Li, J., and Zhu, X. (2020). Leonurine Promotes Cisplatin Sensitivity in Human Cervical Cancer Cells Through Increasing Apoptosis and Inhibiting Drug-Resistant Proteins. Drug Des Devel Ther 14:1885–1895.

Lin, Y., Li, Y., Li, X., Liu, X., Wang, X., Yu, M., Zhu, Y., and Du, M. (2021). SCM-198 ameliorates endometrial inflammation via suppressing the LPS-JNK-cJUN/cFOS-TLR4-NF-κB pathway. Acta biochimica et biophysica Sinica 53:1207–1215.

Liu, Y., Wang, B., Shu, S., Li, Z., Song, C., Liu, D., Niu, Y., Liu, J., Zhang, J., Liu, H., et al. (2021). Analysis of the Coptis chinensis genome reveals the diversification of protoberberine-type alkaloids. Nat Commun 12:3276.

Ma, Y., Cui, G., Chen, T., Ma, X., Wang, R., Jin, B., Yang, J., Kang, L., Tang, J., Lai, C., et al. (2021). Expansion within the CYP71D subfamily drives the heterocyclization of tanshinones synthesis in Salvia miltiorrhiza. Nat Commun 12:685.

Mao, F., Zhang, L., Cai, M.H., Guo, H., and Yuan, H.H. (2015). Leonurine hydrochloride induces apoptosis of H292 lung cancer cell by a mitochondria-dependent pathway. Pharm Biol 53:1684–1690.

Meng, P., Zhu, Q., Yang, H., Liu, D., Lin, X., Liu, J., Fan, J., Liu, X., Su, W., Liu, L., et al. (2019). Leonurine promotes neurite outgrowth and neurotrophic activity by modulating the GR/SGK1 signaling pathway in cultured PC12 cells. Neuroreport 30:247–254.

Miao, L.L., Zhou, Q.M., Peng, C., Liu, Z.H., and Xiong, L. (2019). Leonurus japonicus (Chinese motherwort), an excellent traditional medicine for obstetrical and gynecological diseases: A comprehensive overview. Biomedicine & pharmacotherapy = Biomedecine & pharmacotherapie 117:109060.

Mint Evolutionary Genomics Consortium. Electronic address, b.m.e., and Mint Evolutionary Genomics, C. (2018). Phylogenomic Mining of the Mints Reveals Multiple Mechanisms Contributing to the Evolution of Chemical Diversity in Lamiaceae. Mol Plant 11:1084–1096.

Park, M.N., Um, E.S., Rahman, M.A., Kim, J.W., Park, S.S., Cho, Y., Song, H., Son, S.R., Jang, D.S., Kim, W., et al. (2022). Leonurus japonicus Houttuyn induces reactive oxygen species-mediated apoptosis via regulation of miR-19a-3p/PTEN/PI3K/AKT in U937 and THP-1 cells. Journal of ethnopharmacology 291:115129.

Pitschmann, A., Waschulin, C., Sykora, C., Purevsuren, S., and Glasl, S. (2017). Microscopic and Phytochemical Comparison of the Three Leonurus Species L. cardiaca, L. japonicus, and L. sibiricus. Planta Med 83:1233–1241.

Shang, X., Pan, H., Wang, X., He, H., and Li, M. (2014). Leonurus japonicus Houtt.: ethnopharmacology, phytochemistry and pharmacology of an important traditional Chinese medicine. J Ethnopharmacol 152:14–32.

Srinivasulu, C., Ramgopal, M., Ramanjaneyulu, G., Anuradha, C.M., and Suresh Kumar, C. (2018). Syringic acid (SA) A Review of Its Occurrence, Biosynthesis, Pharmacological and Industrial Importance. Biomed Pharmacother 108:547–557.

Sun, P., Jiao, B., Yang, Y., Shan, L., Li, T., Li, X., Xi, Z., Wang, X., and Liu, J. (2022). WGDI: A user-friendly toolkit for evolutionary analyses of whole-genome duplications and ancestral karyotypes. Molecular plant.

Tian, Z.H., Liu, F., Peng, F., He, Y.L., Shu, H.Z., Lin, S., Chen, J.F., Peng, C., and Xiong, L. (2021). New lignans from the fruits of Leonurus japonicus and their hepatoprotective activities. Bioorganic chemistry 115:105252.

Tu, L., Su, P., Zhang, Z., Gao, L., Wang, J., Hu, T., Zhou, J., Zhang, Y., Zhao, Y., Liu, Y., et al. (2020). Genome of Tripterygium wilfordii and identification of cytochrome P450 involved in triptolide biosynthesis. Nat Commun 11:971.

Wang, J., Mao, Y., Ma, Y., Yang, J., Jin, B., Lin, H., Tang, J., Zeng, W., Zhao, Y., Gao, W., et al. (2022). Diterpene synthases from Leonurus japonicus elucidate epoxy-bridge formation of spiro-labdane diterpenoids. Plant Physiol 189:99–111.

Xiong, L., Peng, C., Zhou, Q.M., Wan, F., Xie, X.F., Guo, L., Li, X.H., He, C.J., and Dai, O. (2013). Chemical composition and antibacterial activity of essential oils from different parts of Leonurus japonicus Houtt. Molecules (Basel, Switzerland) 18:963–973.

Xu, Z., Gao, R., Pu, X., Xu, R., Wang, J., Zheng, S., Zeng, Y., Chen, J., He, C., and Song, J. (2020). Comparative Genome Analysis of Scutellaria baicalensis and Scutellaria barbata Reveals the Evolution of Active Flavonoid Biosynthesis. Genomics Proteomics Bioinformatics 18:230–240.

Yang, B., Hu, Y., Cheng, N., Su, Z., Zhong, Y., Cao, Z., Cao, L., Huang, W., Wang, Z., and Xiao, W. (2020). Anti-inflammatory labdane diterpenoids from Leonurus japonicus Houtt. Phytochemistry 173:112223.

Yue, G.G., Liang, X.X., Li, X.L., Lee, J.K., Gao, S., Kwok, H.F., Lau, C.B., and Xiao, W.L. (2020). Immunomodulatory and antitumour bioactive labdane diterpenoids from Leonurus japonicus. The Journal of pharmacy and pharmacology 72:1657–1665.

Zhan, C., Shen, S., Yang, C., Liu, Z., Fernie, A.R., Graham, I.A., and Luo, J. (2022). Plant metabolic gene clusters in the multi-omics era. Trends Plant Sci 27:981–1001.

Zhang, Q.Y., Wang, Z.J., Miao, L., Wang, Y., Chang, L.L., Guo, W., and Zhu, Y.Z. (2019). Neuroprotective Effect of SCM-198 through Stabilizing Endothelial Cell Function. Oxid Med Cell Longev 2019:7850154.

Zhang, R.H., Liu, Z.K., Yang, D.S., Zhang, X.J., Sun, H.D., and Xiao, W.L. (2018). Phytochemistry and pharmacology of the genus Leonurus: The herb to benefit the mothers and more. Phytochemistry 147:167–183.

Zhang, X.J., Zhong, W.M., Liu, R.X., Wang, Y.M., Luo, T., Zou, Y., Qin, H.Y., Li, X.L., Zhang, R., and Xiao, W.L. (2020). Structurally Diverse Labdane Diterpenoids from Leonurus japonicus and Their Anti-inflammatory Properties in LPS-Induced RAW264.7 Cells. Journal of natural products 83:2545–2558.

Zhao, Q., Yang, J., Cui, M.Y., Liu, J., Fang, Y., Yan, M., Qiu, W., Shang, H., Xu, Z., Yidiresi, R., et al. (2019). The Reference Genome Sequence of Scutellaria baicalensis Provides Insights into the Evolution of Wogonin Biosynthesis. Mol Plant 12:935–950.

Zheng, S., Zhuang, T., Tang, Y., Wu, R., Xu, T., Leng, T., Wang, Y., Lin, Z., and Ji, M. (2021). Leonurine protects against ulcerative colitis by alleviating inflammation and modulating intestinal microflora in mouse models. Exp Ther Med 22:1199.

Zhou, F., Liu, F., Liu, J., He, Y.L., Zhou, Q.M., Guo, L., Peng, C., and Xiong, L. (2020). Stachydrine promotes angiogenesis by regulating the VEGFR2/MEK/ERK and mitochondrial-mediated apoptosis signaling pathways in human umbilical vein endothelial cells. Biomedicine & pharmacotherapy = Biomedecine & pharmacotherapie 131:110724.

Zhu, Y.Z., Wu, W., Zhu, Q., and Liu, X. (2018). Discovery of Leonuri and therapeutical applications: From bench to bedside. Pharmacol Ther 188:26–35.

