## Supplemental data for "Comparative genomic analysis between *Leonurus japonicus* and *Leonurus sibiricus*"

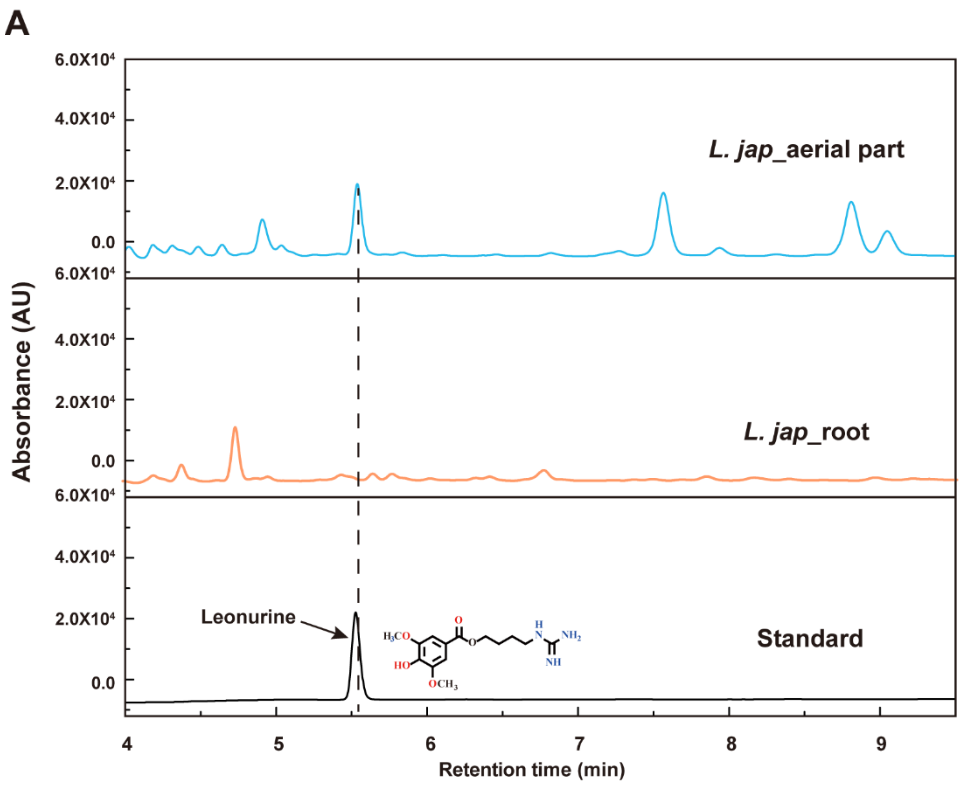


**Figure S1** LC-MS analysis of leonurine contents in leaves and roots of *L. japonicus*.

**A.**

***L. japonicus* (858)**

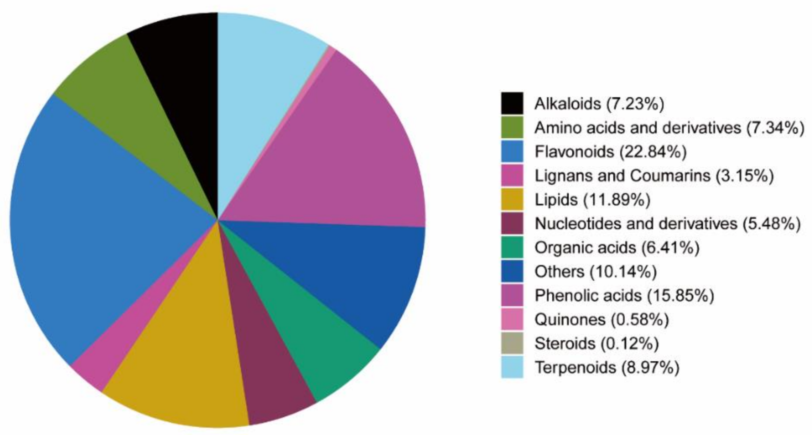


**B.**

***L. sibiricus* (888)**

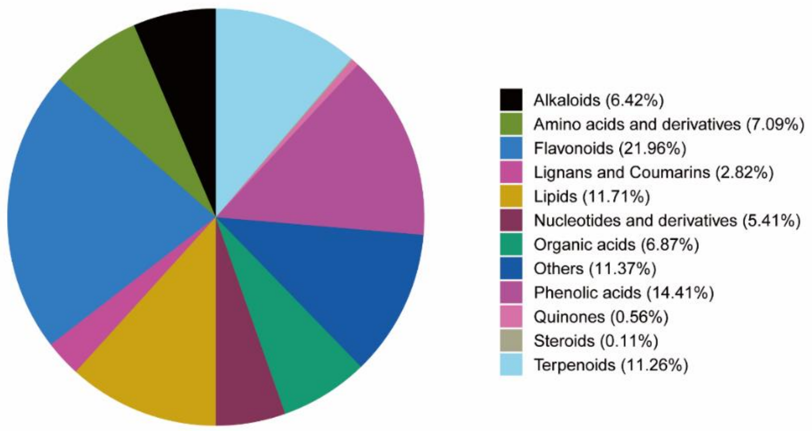


**Figure S2** Classification of secondary metabolites based on the broad-target metabolome in the aerial parts of *L. japonicus* (A) and *L. sibiricus* (B).

**A.**

***L. japonicus***

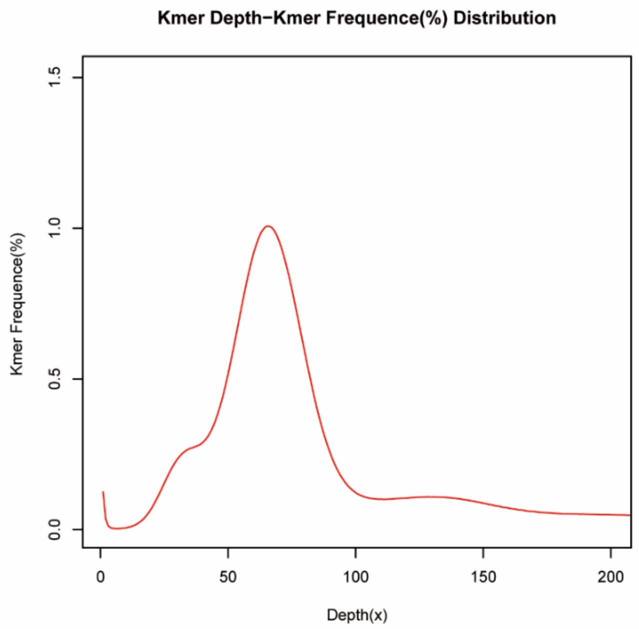


**B.**

***L. sibiricus***

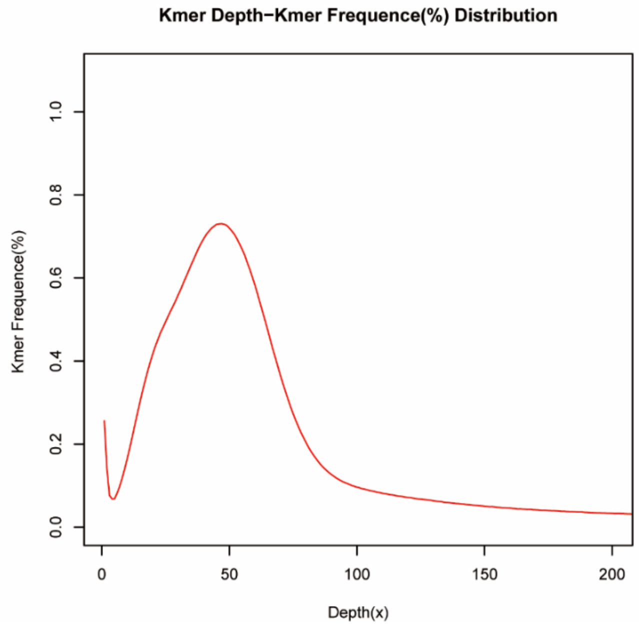


**Figure S3** *K*-mer analysis for estimating the genome size of *L. japonicus* (A) and *L. sibiricus* (B).

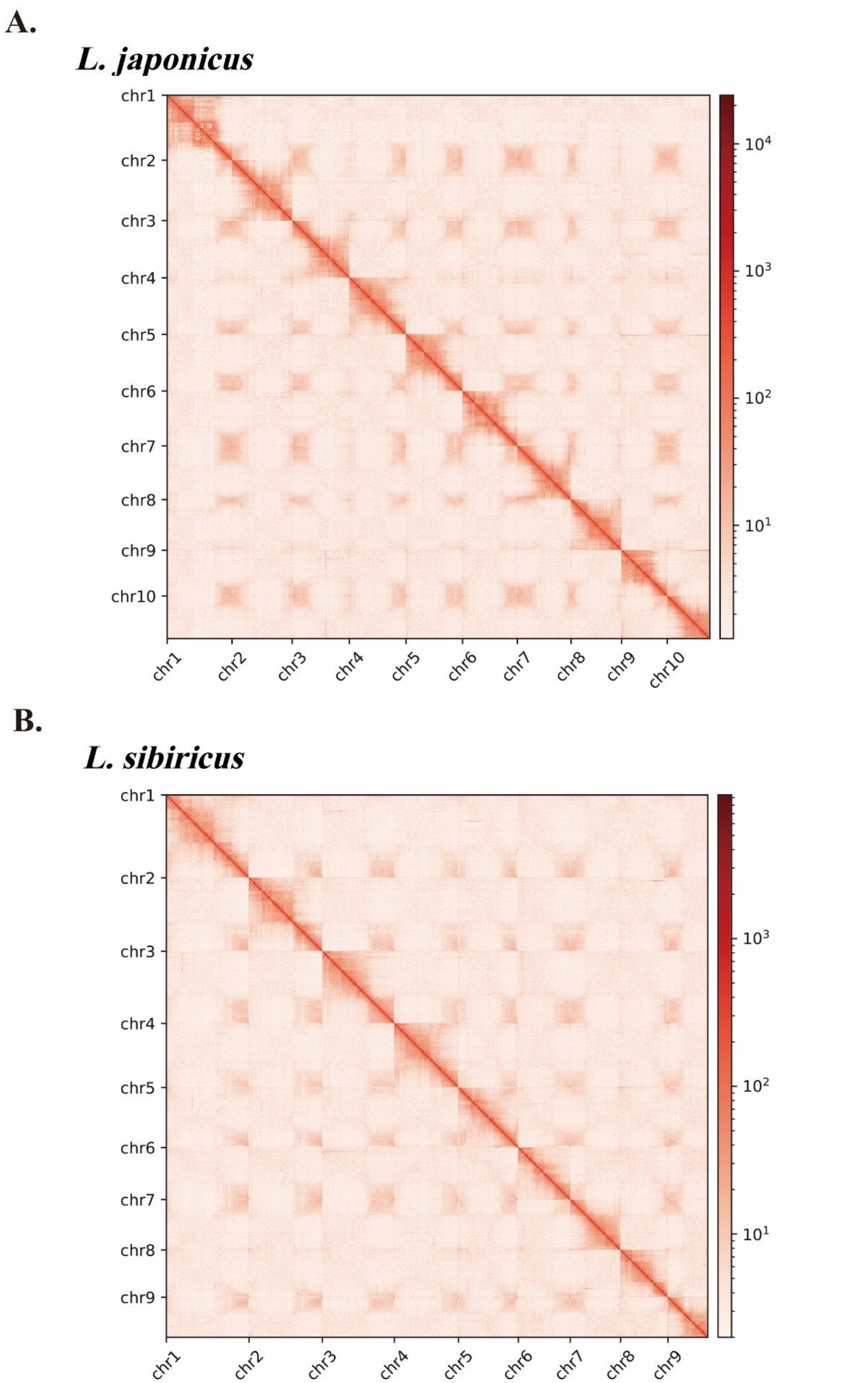


**Figure S4** Intensity signal heatmap of Hi-C chromosome interactions of *L. japonicus* and *L. sibiricus*.

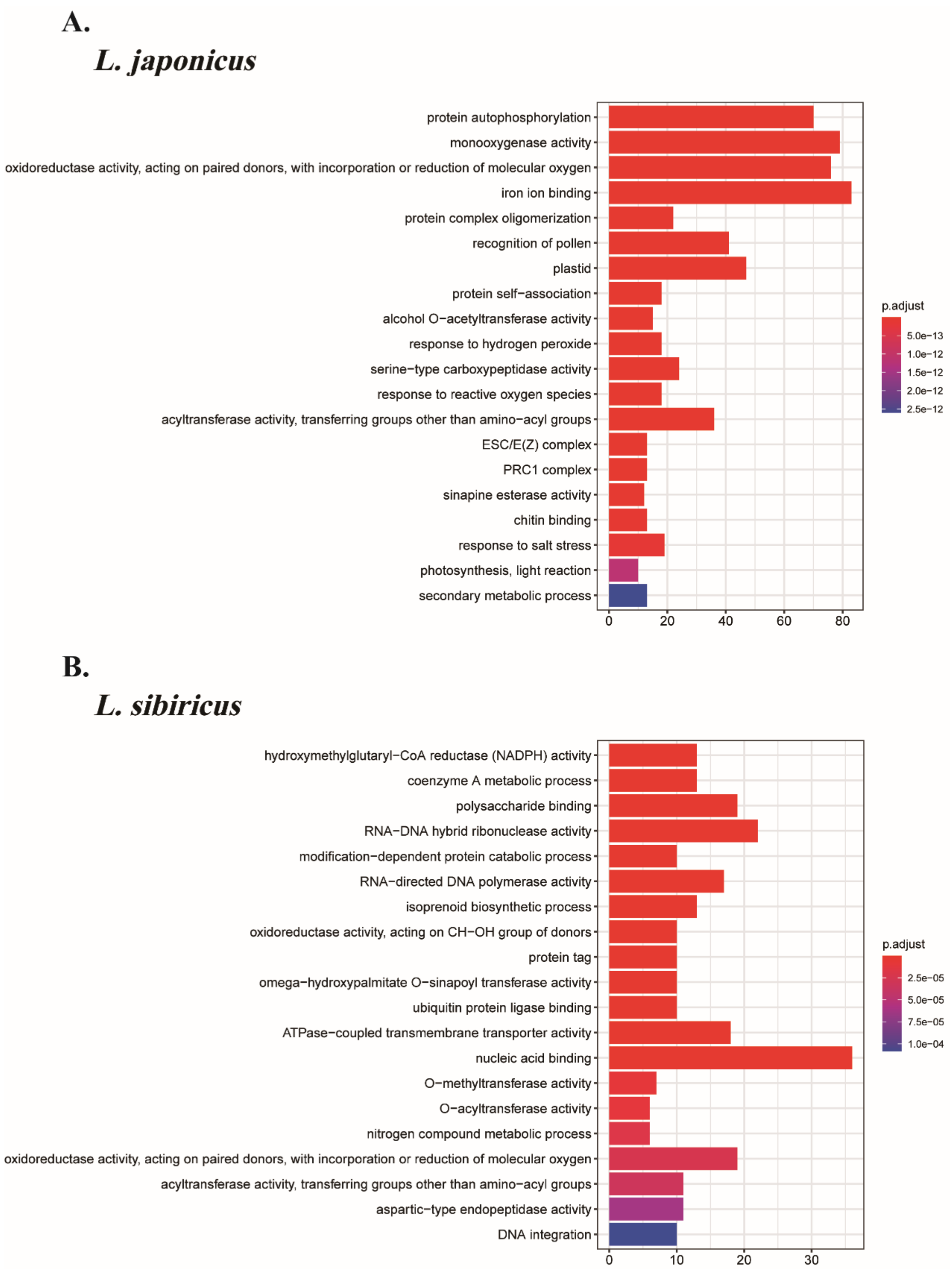


**Figure S5** GO enrichment results of the expanded gene families in *L. japonicus* and *L. sibiricus* (top 20 for display).

**Table S1** Survey statistic results of *L. japonicus*.

| Sample | K-mer | K-mer Number | K-mer Depth | Genome Size (M) | Data Size (G) | X | Heterozygous Ratio (%) | Duplication Ratio (%) |
|---|---|---|---|---|---|---|---|---|
| *L. jap* | 19 | 35,367,187,460 | 63.75 | 553.75 | 55.75 | 100.68 | 0.73 | 58.39 |

**Table S2** Survey statistical results of *L. sibiricus*.

| Sample | K-mer | K-mer Number | K-mer Depth | Genome Size (M) | Data Size | X | Heterozygous Ratio (%) | Duplication Ratio (%) |
|---|---|---|---|---|---|---|---|---|
| *L. sib* | 19 | 23,040,594,040 | 49.8 | 465.78 | 34.14 | 73.3 | 1.91 | 66.39 |

**Table S3** Summary of repeat contents in the *L. japonicus* genome.

| Type | Length (bp) | % in genome | Length (bp) | % in genome | Length (bp) | % in genome |
|---|---|---|---|---|---|---|
| DNA | 4,191,381 | 0.81 | 29,450,251 | 5.68 | 30,250,963 | 5.84 |
| LINE | 4,147,271 | 0.80 | 9,441,898 | 1.82 | 10,282,676 | 1.98 |
| SINE | 0 | 0.00 | 8,443 | 0.00 | 8,443 | 0.00 |
| LTR | 68,059,721 | 13.13 | 211,579,519 | 40.83 | 212,996,860 | 41.10 |
| LTR-Gypsy | 20,880,805 | 4.03 | 85,385,156 | 16.48 | 86,184,128 | 16.63 |
| LTR-Copia | 46,572,988 | 8.99 | 124,544,051 | 24.03 | 125,049,653 | 24.13 |
| Satellite | 0 | 0.00 | 734,062 | 0.14 | 734,062 | 0.14 |
| Simple_repeat | 0 | 0.00 | 349,492 | 0.07 | 349,492 | 0.07 |
| Other | 0 | 0.00 | 6,415 | 0.00 | 6,415 | 0.00 |
| Unknown | 24,375 | 0.00 | 86,528,278 | 16.70 | 86,552,636 | 16.70 |
| Total | 76,416,034 | 14.75 | 330,036,531 | 63.69 | 344,639,335 | 66.51 |

**Table S4** Summary of Repeat contents in the *L. sibiricus* genome.

| Type | Length (Bp) | % in genome | Length (bp) | % in genome | Length (bp) | % in genome |
|---|---|---|---|---|---|---|
| DNA | 3,579,907 | 0.76 | 28,388,455 | 6.01 | 29,147,689 | 6.17 |
| LINE | 3,708,316 | 0.79 | 9,407,344 | 1.99 | 10,171,960 | 2.15 |
| SINE | 0 | 0.00 | 6,995 | 0.00 | 6,995 | 0.00 |
| LTR | 64,040,923 | 13.56 | 190,856,438 | 40.41 | 192,098,379 | 40.67 |
| LTR-Gypsy | 17,291,402 | 3.66 | 62,193,868 | 13.17 | 62,801,078 | 13.30 |
| LTR-Copia | 46,379,892 | 9.82 | 127,597,343 | 27.02 | 128,112,427 | 27.13 |
| Satellite | 0 | 0.00 | 665,888 | 0.14 | 665,888 | 0.14 |
| Simple_repeat | 0 | 0.00 | 303,570 | 0.06 | 303,570 | 0.06 |
| Other | 0 | 0.00 | 3,452 | 0.00 | 3,452 | 0.00 |
| Unknown | 18,060 | 0.00 | 79,181,289 | 16.77 | 79,199,349 | 16.77 |
| Total | 71,343,538 | 15.11 | 301,574,907 | 63.85 | 308,846,385 | 65.39 |

**Table S5** The statistical results of noncoding RNA of the *L. japonicus* genome.

| Type | | Copy | Average length (bp) | Total length (bp) | % of genome |
|---|---|---|---|---|---|
| miRNA | | 107 | 124 | 13,301 | 0.002567 |
| tRNA | | 585 | 75 | 44,153 | 0.008520 |
| rRNA | rRNA | 238 | 289 | 68,717 | 0.013261 |
| | 18S | 26 | 1520 | 39,513 | 0.007625 |
| | 28S | 41 | 198 | 8,100 | 0.001563 |
| | 5.8S | 28 | 155 | 4,333 | 0.000836 |
| | 5S | 143 | 117 | 16,771 | 0.003236 |
| snRNA | snRNA | 1,704 | 109 | 185,433 | 0.035784 |
| | CD-box | 1,516 | 106 | 160,133 | 0.030902 |
| | HACA-box | 54 | 114 | 6,134 | 0.001184 |
| | splicing | 134 | 143 | 19,166 | 0.003699 |
| | scaRNA | 0 | 0 | 0 | 0.000000 |

**Table S6** The statistical results of noncoding RNA of the *L. sibiricus* genome.

| Type | | Copy | Average length (bp) | Total length (bp) | % of genome |
|---|---|---|---|---|---|
| miRNA | | 95 | 122 | 11,624 | 0.002461 |
| tRNA | | 524 | 75 | 39,235 | 0.008307 |
| rRNA | rRNA | 219 | 252 | 55,170 | 0.011681 |
| | 18S | 34 | 918 | 31,227 | 0.006612 |
| | 28S | 23 | 188 | 4,331 | 0.000917 |
| | 5.8S | 14 | 158 | 2,218 | 0.000470 |
| | 5S | 148 | 118 | 17,394 | 0.003683 |
| snRNA | snRNA | 1,449 | 109 | 158,587 | 0.033578 |
| | CD-box | 1,276 | 106 | 134,740 | 0.028529 |
| | HACA-box | 49 | 115 | 5,630 | 0.001192 |
| | splicing | 124 | 147 | 18,217 | 0.003857 |
| | scaRNA | 0 | 0 | 0 | 0.000000 |

**Table S7** Statistics of colinearity gene pairs between *L. japonicus* and *L. sibiricus*.

| *L.jap* / *L.sib* | **chr1** | **chr2** | **chr3** | **chr4** | **chr5** | **chr6** | **chr7** | **chr8** | **chr9** | **chr10** |
|---|---|---|---|---|---|---|---|---|---|---|
| **chr1** | 13 | 4 | 5 | 22 | 1209 | 1602 | 70 | 16 | 36 | 0 |
| **chr2** | 2009 | 810 | 0 | 31 | 15 | 5 | 0 | 0 | 0 | 30 |
| **chr3** | 0 | 834 | 860 | 24 | 821 | 288 | 17 | 17 | 79 | 12 |
| **chr4** | 0 | 411 | 1537 | 16 | 8 | 0 | 11 | 0 | 10 | 5 |
| **chr5** | 6 | 0 | 31 | 17 | 42 | 14 | 2719 | 51 | 0 | 83 |
| **chr6** | 0 | 15 | 5 | 1636 | 25 | 21 | 7 | 35 | 0 | 52 |
| **chr7** | 27 | 0 | 5 | 0 | 12 | 0 | 154 | 42 | 19 | 1546 |
| **chr8** | 0 | 9 | 0 | 49 | 0 | 13 | 0 | 932 | 5 | 0 |
| **chr9** | 0 | 44 | 0 | 0 | 8 | 56 | 0 | 0 | 1533 | 9 |
